# Excitation Spectral Phasor Microscopy (ExSPM) unveils spatiotemporal heterogeneities in polarity of intracellular lipid droplets

**DOI:** 10.1101/2025.04.03.647137

**Authors:** Jiayi Liu, Mingxuan Tang, Rui Yan, Jinhong Yan, Yi He, Ruirong Wang, Zhipeng Zhang, Jiankai Xia, Igor Kireev, Ke Xu, Kun Chen

## Abstract

Spatiotemporal heterogeneities in intracellular LD polarity are central to cellular homeostasis but remain largely unexplored. Here, we introduce Excitation Spectral Phasor Microscopy (ExSPM), which integrates high-throughput excitation spectral imaging with phasor analysis to map LD polarity with exceptional sensitivity and spatiotemporal resolution. Using the lipophilic dye Nile Red, ExSPM revealed extensive heterogeneity and dynamics of LD polarity within single cells, reflecting lipid metabolism and energy status. Notably, high-polarity LDs emerged dynamically at lysosome- and mitochondria-contact sites, revealing pathway-specific kinetics of LD catabolism that distinguish lipophagy from neutral lipolysis. ExSPM further uncovered a tightly interconnected regulatory network among lipolytic pathways, in which perturbation of one pathway reshapes others while preserving overall energy supply. ExSPM thus provides a versatile platform to dissect intracellular compositional heterogeneity and the coordinated regulation of lipid metabolism, offering new insights into organelle function and cellular homeostasis.

## Introduction

Lipid droplets (LDs) are dynamic organelles essential for cellular lipid storage and metabolism^1^. Beyond serving as reservoirs of neutral lipids, they actively interact with other organelles to regulate energy homeostasis and lipid signaling^2, 3^. Structurally, LDs consist of a core of nonpolar neutral lipids, such as triacylglycerol (TAG) and cholesterol ester (CE), surrounded by a more polar phospholipid monolayer embedded with specialized surface proteins^4^. Although once considered compositionally uniform, LDs are now known to be highly heterogeneous, both across different cell types and within individual cells^5, 6^. This heterogeneity reflects varying metabolic demands and is increasingly recognized as important in cellular regulation and disease^7^. However, most studies have focused on differences in the surface monolayer—its protein content and phospholipid composition—revealing roles in protein recruitment, lipid metabolism, and interactions with other organelles^6, 8–11^. In contrast, the composition of the neutral lipid core—critical to LD metabolic function—remains poorly characterized, particularly at the single-cell level. A key factor underlying this core compositional heterogeneity is LD polarity, which reflects differences in the internal chemical environment^12^. Polarity influences how lipids are packed, how lipid phases separate, and how proteins interact with the LD surface^13, 14^. These polarity-dependent variations shape the functional diversity of LDs and underscore their active and adaptable role in cell physiology.

Fluorescence solvatochromic dyes, such as Nile Red and its derivatives, offer a powerful means to probe LD polarity due to their red-shifted fluorescence in polar environments^15–17^. Recent studies employing these dyes have revealed significant polarity differences among normal and cancer cells, various cancer types, muscle-damaged cells, and liver lesion cells^13, 14, 18, 19^. These findings offer new insights into LD composition and function while highlighting their potential as diagnostic markers for disease detection. Despite growing recognition of polarity differences between cell types, how LD polarity varies within individual cells remains poorly understood—yet this intracellular heterogeneity is likely crucial for metabolic adaptation and cell function. Even within a single cell, LDs can exhibit distinct polarity profiles influenced by their spatial context, such as proximity to specific organelles, local metabolic activity, developmental stage, or transient stressors like oxidative hotspots and nutrient fluctuations. For example, LDs closely associated with mitochondria have been shown to undergo liquid–crystalline phase transitions^20^ and facilitate the transfer of harmful proteins to mitigate mitochondrial damage^21^—both processes are likely to drive the changes in polarity. Thus, deciphering spatiotemporal variations in intracellular LD polarity is essential for understanding their morphological, chemical, and compositional heterogeneities, as well as the dynamic metabolic microenvironments that shape cellular behavior and disease progression. Although a recent study using newly developed polarity-sensitive probes reported intracellular LD polarity differences^17^, the sensitivity and temporal resolution were insufficient to resolve the underlying mechanisms.

Addressing this challenge requires advanced imaging techniques capable of resolving subtle LD polarity variations with high spatial and temporal resolutions. Current approaches using solvatochromic dyes primarily rely on ratiometric fluorescence measurements with two excitation or emission channels^14, 16, 17, 19, 22^, limiting their sensitivity to subtle intracellular differences. While point-scanning spectral imaging can capture full emission spectra to enhance sensitivity to spectral variations^15^, it imposes substantial constraints on temporal resolution. Unlike emission-based methods, we previously developed high-throughput excitation spectral microscopy (ExSM), which rapidly scans frame-synchronized excitation wavelengths for the entire imaging field to acquire wavelength-resolved images^23–25^. This approach enables spectral unmixing to quantify fluorophore abundances, facilitating novel imaging strategies for bi-state biosensors. However, spectral unmixing cannot disentangle multi-state biosensors, such as polarity-sensitive dyes with full-color solvatochromic fluorescence, as their spectra shift continuously rather than consisting of discrete states.

To overcome these limitations, we here developed <u>Ex</u>citation <u>S</u>pectral <u>P</u>hasor <u>M</u>icroscopy (ExSPM), a technique that integrates high-throughput excitation fluorescence spectral microscopy with phasor analysis. ExSPM detects subtle spectral responses of fluorophores to local intracellular environments without prior spectral knowledge, enabling the direct visualization of intracellular spatiotemporal heterogeneities. Using the lipophilic fluorophore Nile Red, ExSPM revealed rich heterogeneities and dynamics in LD polarity, shaped by lipid metabolism and cellular energy homeostasis. It further quantified pathway-specific contributions to lipid droplet catabolism, exposing a tightly interconnected regulatory network of lipolytic pathways. These findings underscore ExSPM’s transformative potential to advance our understanding of spatiotemporal heterogeneities in intracellular composition and function.

## Results

Spectral phasor analysis is a non-fitting, Fourier-based method for extracting information from fluorescence lifetime^26^, and more recently fluorescence spectral imaging^27–30^. By converting high-dimensional spectral data into a 2D phasor space, it enables visualization of pixel ensembles, revealing population distributions and interpixel correlations inaccessible to fitting analysis^31–34^. Widely used in multiplexed fluorescence microscopy, it excels in spectral unmixing, compression, denoising, and computational efficiency^28^. Here, we repurpose phasor analysis for excitation spectral microscopy (ExSPM) to start assessing its multiplexing performance in five-color intracellular imaging. Using eight excitation wavelengths based on our previously reported ExSM setup^23, 25^, spectrally overlapping fluorophores labeling five distinct subcellular targets were successfully unmixed with minimal crosstalk by leveraging linear additivity in phasor space (Fig. S1 and the Supplementary Note 1 for details). While spectral phasor analysis has been widely used for multi-color imaging, it has also been applied to characterize spectral variations of the membrane probe LAURDAN^34–36^. Here, we extend this capability by demonstrating that ExSPM offers exceptional sensitivity in detecting subtle excitation spectral shifts, thereby unveiling spatiotemporal heterogeneities that governs composition and function of LDs.

### ExSPM unveils distinct compositional polarity heterogeneity of LDs within cells

Intracellular LDs are highly dynamic organelles central to lipid metabolism, membrane synthesis, and lipid-derived signaling. We started with Nile Red, a solvatochromic lipid dye commonly used to report changes in the chemical polarity of lipids in cells^15, 37–41^. Previous studies based on emission spectral shifts showed that mitochondrial and endoplasmic reticulum (ER) membranes are more polar and structurally less ordered than the plasma membrane, exhibiting red-shifted fluorescence^38^. In this study, we examined the excitation spectra of Nile Red on different membranes in live COS-7 cells (Supplementary Fig. S2). Cells were imaged in a buffer containing a high concentration of Nile Red (3.3 μM), ensuring that the cell plasma membrane and the membranes of intracellular organelles were well labeled and visualized. Consistent with prior findings, mitochondrial and ER membranes showed redder shifts relative to the plasma membrane, while LDs exhibited the most blue-shifted excitation spectra, indicating lower polarity and a more ordered phase. The excitation spectra of Nile Red dissolved in solvents of various known polarities also demonstrate the spectral shifts depending on solvent polarity (Supplementary Fig. S2c). Interestingly, some LDs also exhibited unexpected red-shift spectra (the magenta arrow in Supplementary Fig. S2).

To understand the substantial polarity heterogeneities of individual LDs, we then utilized ExSPM to examine potential changes in the underlying excitation spectra of Nile Red at high spectral and spatial resolutions. At a low concentration (∼30 nM), Nile Red preferentially stained LDs over cell membranes (Fig. 1a and Supplementary Fig. S3), minimizing interference from other structures. The phasor plot of the spectral images grouped pixels with similar spectral profiles into clusters, enabling direct visual inspection of potential spectral shifts without prior spectral knowledge (Fig. 1b, where *G* and *S* are the real and imaginary coefficients). Unexpectedly, the phasor plot exhibited a broad distribution in *G*-*S* space, suggesting spatial heterogeneities within LDs, with a subset in the lower-right plane potentially indicating high polarity. Revisiting the excitation spectra of six LDs within the cell (marked with arrows in Fig. 1a), we found notable differences, including a striking ∼60 nm blue shift from LD1 to LD6 (Fig. 1c).

**Figure 1.**
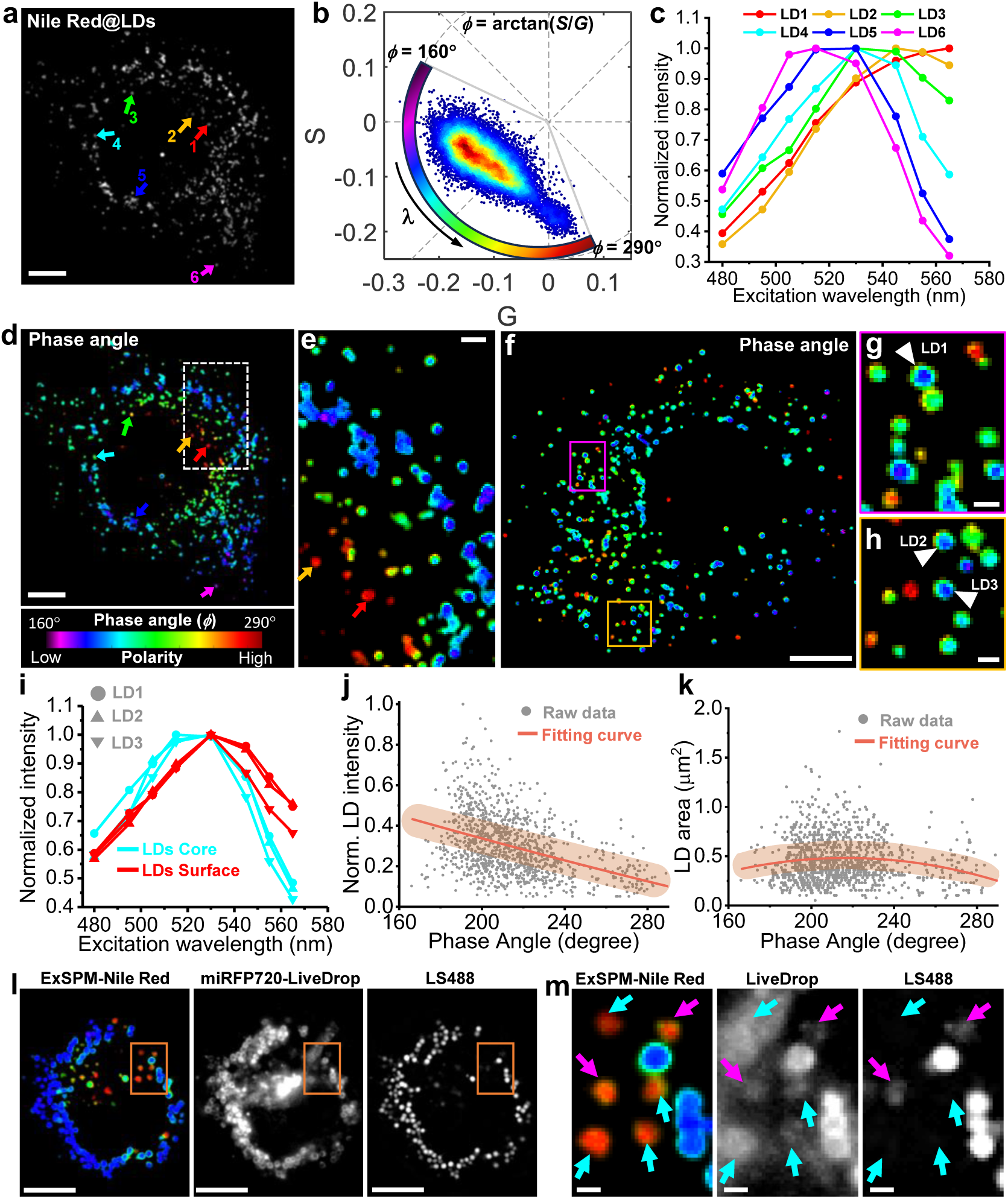
ExSPM unveils distinct compositional polarity heterogeneity of LDs within cells. **a.** Conventional fluorescence image of LDs stained with Nile Red at a low concentration (30 nM) in a COS-7 cell. **b.** ExSPM represents excitation spectral data as a phasor plot, a 2D histogram of the real (*G*) and imaginary (*S*) Fourier components at a single harmonic. **c.** Excitation spectra of six individual LDs indicated by arrows in a. **d.** High-resolution polarity mapping of intracellular LDs, color-coded by the phase angle derived from the phasor plot in b. **e.** Magnified view of the white-boxed subregion in d. **f.** An LD polarity map of another cell obtained with ExSPM. **g, h.** Zoomed-in views of the boxed regions in f. **i.** Spectral differences between the core and surface of three LDs marked by arrowheads in. **j.** Distributions of the measured phase angles and fluorescence intensities for single LDs. **k.** Distributions of the measured phase angles and areas for single LDs. Points, results from individual LDs. **l.** Live-cell concurrent imaging of a COS-7 cell showing the ExSPM-derived LD polarity map (Nile Red staining), miRFP720-tagged LiveDrop, and LipidSpot488 (LS488)-labeled LDs. **m.** Zoomed-in views of the boxed regions in l. Cyan arrows: high-polarity LDs marked by LiveDrop only; Magenta arrows: High-polarity LDs marked by both LS488 and LiveDrop. Scale bars: 10 μm (a, d, f, l); 2 μm (e); 1 μm (g, h, m).

To present both spectral and spatial information for each LD, we calculated the phase angle of each pixel and used this value to map it onto a continuous color scale (160° – 290°). Physically, the phase angle (*ϕ*=arctan(S/G)) is linear with the mean wavelength (the center of mass of the excitation spectrum) from spectral phasor transformation (Fig. 1c and Supplementary Note 1). Accordingly, a small phase angle denotes blue-shift spectrum corresponding to low polarity and ordered phases in LDs. The resultant “true-color” images revealed striking spectral differences among LDs (Fig. 1c and d, and Supplementary Fig. S3 and S4) with a strong phase shift up to 130°. This extensive dynamic range, coupled with ExSPM’s high-throughput spectral detection, enables the identification of LD subpopulations with distinct polarities, allowing for highly sensitive mapping of LD compositional and functional heterogeneities across entire cells at high spatiotemporal resolutions. In contrast, standard ratiometric imaging lacks the sensitivity to reliably detect this level of heterogeneity (Supplementary Fig. S5).

ExSPM revealed distinct spatial patterns of LD polarity. In some cells, red-shifted LDs were dispersed throughout the cytoplasm (Fig. 1f; Supplementary Fig. S4–S6), but in most cases they accumulated in the perinuclear region (Fig. 1d-e; Supplementary Fig. S3 and S6). The proportion of red-shifted LDs varied widely between cells (Supplementary Fig. S6), indicating substantial heterogeneity. Within individual LDs, polarity often varied from core to surface (Fig. 1g–i; Supplementary Fig. S3-S4). This gradient is consistent with less-polar TAG/CE-rich cores compared to phospholipid monolayers, and may also reflect diffraction-related signal loss at droplet edges^16^. Quantitative analysis showed that low-polarity droplets (smaller phase angles) were brighter, while red-shifted droplets were dimmer (Fig. 1j; Supplementary Fig. S3), in line with Nile Red’s higher quantum yield in nonpolar environments. By contrast, polarity showed little dependence on droplet size (Fig. 1k; Supplementary Fig. S4 and S7), a result further validated in artificial adiposomes of varying diameters (Supplementary Fig. S8). These measurements were robust across Nile Red concentrations, both in cells and in pure TAG droplets (Supplementary Fig. S9 and S10). Importantly, ExSPM demonstrated high-precision polarity mapping (∼1.22°), which enabled clear discrimination between TAG and CE droplets (Supplementary Fig. S11).

To confirm that the observed polarity heterogeneities originate from bona fide LDs rather than other lipid-rich, high-polarity organelles, we used the acid-stable LD dye LipidSpot488 (LS488, Supplementary Fig. S12) together with established LD markers LiveDrop^42, 43^ and PLIN2^6^, both tagged with the pH-insensitive fluorescent protein mCherry, for colocalization analysis. Dual-color imaging of mCherry-LiveDrop and LS488 in live COS-7 cells revealed that all LS488-positive droplets were encompassed by LiveDrop signals (Supplementary Fig. S12), a result reproduced with mCherry-PLIN2 and LS488 (Supplementary Fig. S13). These findings demonstrate the high specificity of LS488 for LDs through selective binding to neutral lipid-rich cores. While PLIN2 puncta rarely lacked LS488 staining, a subset of LiveDrop-positive structures frequently did (cyan arrows in Fig. S12), likely representing LDs depleted of neutral lipids while retaining mCherry-LiveDrop signal. Notably, starved cells with extensive lipid consumption showed increased abundance of isolated LiveDrop puncta. (Supplementary Fig. S14).

We further performed concurrent imaging of LS488, LiveDrop, and ExSPM-Nile Red. To avoid spectral crosstalk from Nile Red’s broad solvatochromic response, LiveDrop was labeled with miRFP720—the most red-shifted monomeric near-infrared fluorescent protein, which remains fluorescence under acidic conditions as mCherry^44^. A sequential imaging workflow was applied: ExSPM imaging of Nile Red was conducted in live cells first, followed by excitation of miRFP720 at 647 nm, and finally LS488 staining and imaging at 450 nm in the same cells (Fig. 1l-m, Supplementary Fig. S15). This strategy avoids spectral crosstalks among these fluorophores and enables unambiguous LD identification together with quantitative polarity mapping. Most Nile Red–labeled droplets colocalized with both LS488and LiveDrop, and displayed a broad range of polarities, providing compelling evidence for polarity heterogeneity among LDs within individual cells (Fig. 1l-m). A subset of high-polarity LDs colocalized with LiveDrop but lacked LS488 staining, corresponding to droplets with depleted neutral lipid content identified above. Moreover, analysis of LS488 intensity relative to LD polarity showed that increasing polarity was consistently associated with decreasing LS488 signal (Supplementary Fig. S16), indicating that the polarity gradient likely reflects progressive neutral lipid consumption within cells (see more discussions below).

### ExSPM unveils the spatiotemporal heterogeneities in polarity of LDs in live cells

We next harnessed our above capability to map the spatiotemporal dynamics of LD polarity in live cells. Through continuous recording under our 8-wavelength excitation scheme at 10 fps and performing phasor analysis for every pixel, we obtained full-frame LD polarity images at 0.8 s time resolution (Fig. 2a-c, Supplementary Fig. S17, Video 1 and 2). This approach revealed diverse polarity dynamics among individual LDs within a single cell. LDs, as highly dynamic organelles, frequently undergo fission and fusion events critical for lipid biogenesis and catabolism. During LD–LD fusion, we observed a gradual polarity decrease (Fig. 2d-e, Supplementary Fig. S17 and Video 3), whereas fission led to increased polarity (Fig. 2f-g, Supplementary Fig. S17 and Video 4). These changes likely reflect dynamic molecular reorganization within the LD core: fusion may facilitate mixing and homogenization of internal lipid constituents, thereby reducing polarity, whereas fission may result in asymmetric partitioning or localized enrichment of more polar molecules. Even within individual LDs, polarity fluctuated significantly (230° – 260°) over 1-2 minutes (Supplementary Fig. S17e-g), indicating ongoing compositional changes. Notably, fixation with paraformaldehyde and glutaraldehyde preserved the polarity characteristics and heterogeneity of LDs observed in live cells (Supplementary Fig. S18), suggesting that LD compositions remain stable following chemical fixation.

**Figure 2.**
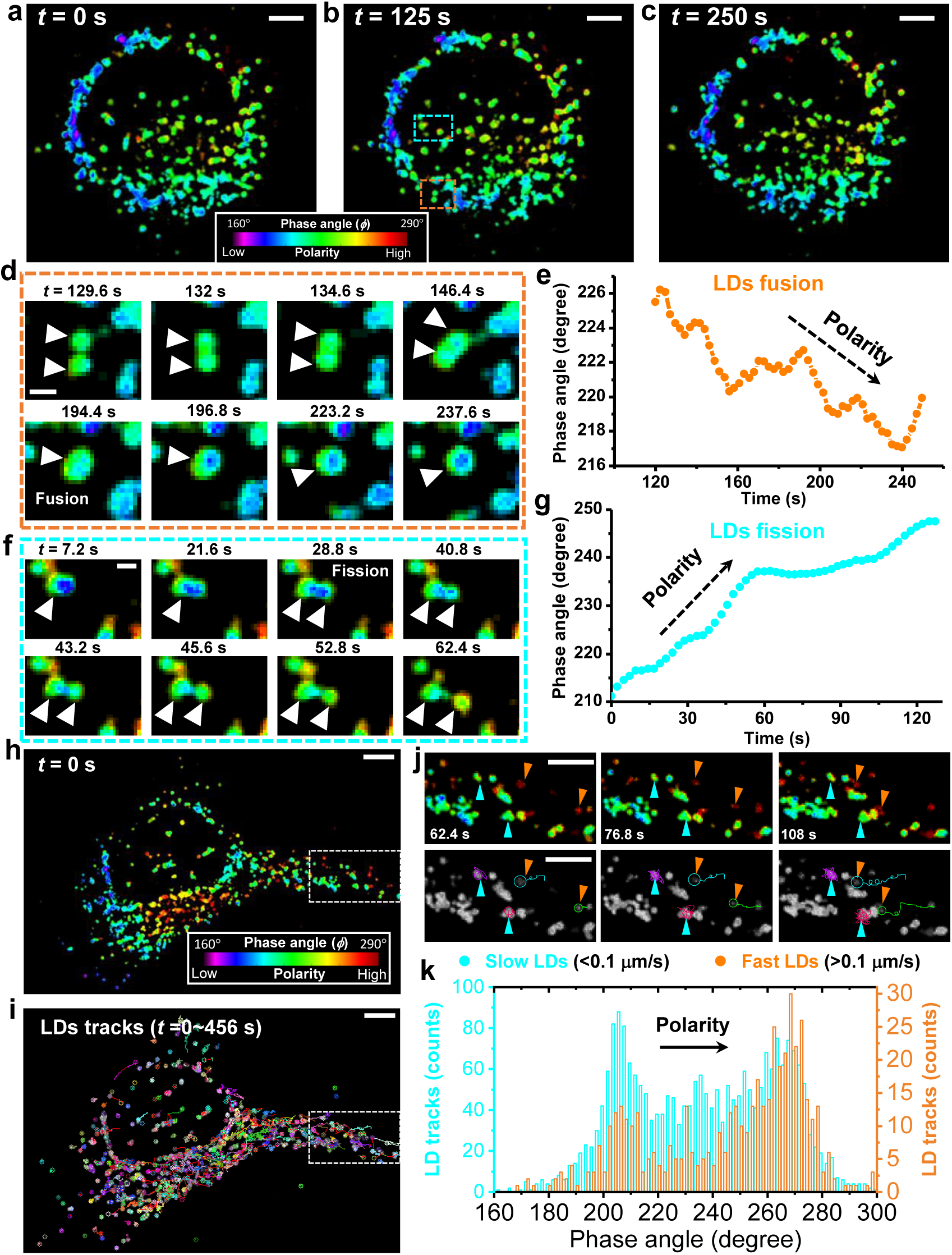
ExSPM unveils the spatiotemporal heterogeneities in polarity of LDs in live cells. **a-c**. ExSPM-derived LD polarity maps of a live COS-7 cell at different time points. **d**. Time sequences for the orange boxed region in b, showing LD fusion and associated polarity changes. **e.** Temporal variation in polarity for LDs indicated by arrowheads in d, highlighting a gradual polarity decrease during LD fusion. **f**. Time-lapse sequence of the cyan-boxed region in b, showing LD fission and associated polarity changes. **g.** Temporal variation in polarity for LDs indicated by arrowheads in f, highlighting a gradual polarity increase during LD fission. **h**. ExSPM-derived LD polarity map of another live COS-7 cell at *t* =0 s. **i**. Detected LDs and their local trajectories accumulated over 0– 456 s, indicated by different colors. The movement of LDs were automatically tracked using a particle tracker model (MosaicSuite, see Supplementary Video 5). **j**. Zoomed-in views of the boxed regions in h and i. Top: ExSPM-derived LD polarity map. Bottom: LDs tracks. Fast and high-polarity LDs are marked with orange arrowheads. Slow and low-polarity LDs are marked with cyan arrowheads. **k**. Distribution of measured phase angles (polarity) for fast (orange) and slow (cyan) LDs, classified based on a speed threshold of 0.1 μm/s. The experiments above were performed with 8-wavelength excitation cycles at 10fps. Each phasor image was generated by averaging three consecutive 8-frame sequences, resulting in an effective acquisition time of 2.4 s. Scale bar: 10 μm (a-c, h, i); 1 μm (d, f); 5 μm (j).

LDs exhibit significant mobility (Fig. 2h, Supplementary Video 5), with slow movements appearing random and diffusional, while fast movements are more directional (Fig. 2i and 2j). Using the particle tracker model (MosaicSuite)^45^, we tracked individual LDs (color-coded in Fig. 2i) and quantified their velocities. One-step movement analysis of each LDs (Fig. 2k) revealed that high-polarity LDs tend to move faster (orange arrows in Fig. 2j), while low-polarity LDs predominantly move more slowly (cyan arrows in Fig. 2j), a correlation evident in Supplementary Videos 5 and 6. The rapid, directional motion of high-polarity LDs likely reflects their association with microtubules, which mediate LD transport for various cellular functions^46^. In contrast, the slower, random movement of low-polarity LDs suggests they diffuse freely within the cytoplasm. An additional hypothesis is lysosome-LD interactions during lipophagy may enhance LD mobility, as lysosomes have been reported to move at twice the speed of LDs^47^ (see more discussions below).

### ExSPM unveils that LDs polarity is closely linked to cellular energy metabolism

We next asked whether LD polarity is influenced by energy metabolism in live cells. Cells balance energy demands by activating fatty acid oxidation during starvation and promoting lipogenesis under nutrient-rich conditions. To test this, COS-7 cells were cultured in full-serum medium, serum-starved medium, or oleic acid (OA)-supplemented medium (Fig. 3a-l). Starvation induced a clear increase in LD polarity, reflected by a red-shifted spectral phase angle compared with full-serum conditions (Fig. 3e vs. Fig. 3f). Statistical analysis further validated this trend (Fig. 3j), consistent with our earlier finding that high-polarity LDs are more mobile (Fig. 2k) and with Raman imaging studies reporting more fast-moving LDs under starvation^48^. Conversely, excess OA led to increased LD accumulation and significantly reduced polarity, both in full-serum (Fig. 3c, g, and k) and FBS-starvation medium (Fig. 3d, h, and l), with stronger effects at higher OA concentrations or longer treatments (Supplementary Fig. S19c-h and Fig. S20). Similar polarity changes were observed in HeLa and A549 cells (Fig. 3m), though HeLa consistently exhibited higher LD polarity than COS-7 and A549 under all conditions (Fig. 3m, and Supplementary Fig. S21-S22).

**Figure 3.**
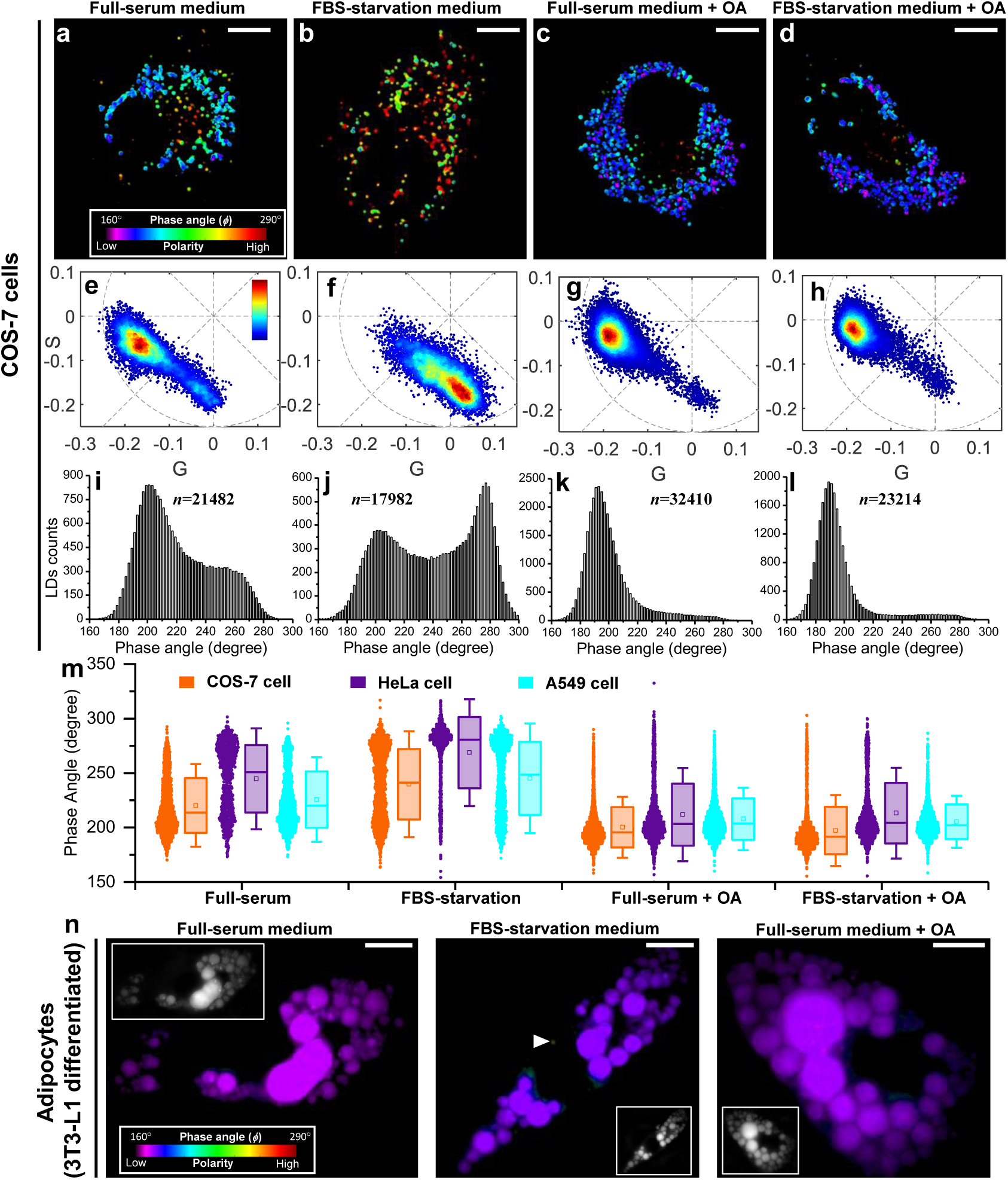
Assessment of LD polarity under different nutrient conditions using ExSPM. **a-d.** ExSPM-derived LD polarity maps of COS-7 cells cultured in full-serum medium (a), FBS-starvation medium (b), and 300 μM OA-supplemented media for 16h (c, d). **e-h.** Corresponding Phasor plot of the same excitation spectra data in a–d, providing a direct visualization of spectral shifts. **i-l.** Distributions of phase angles (polarity) for tens of thousands of LDs across 80-100 cells under the nutrient conditions in a–d. **m.** Statistical analysis of phase angles (polarity) across COS-7, HeLa, and A549 cells under different nutrient conditions. Points, results from individual LDs. **n.** ExSPM-derived LD polarity maps of 3T3-L1 differentiated adipocytes cultured in full-serum medium, FBS-starvation medium, and 300 μM OA-supplemented media for 16h. Inset: fluorescence intensity images. All experiments in each condition were independently repeated at least three times with similar results. Scale bars:10 μm (a-d, n).

These results suggest that nutrient deprivation promotes neutral lipid hydrolysis^49^, releasing progressively more polar metabolites—diacylglycerols, monoacylglycerols, free fatty acids (FFAs), cholesterol, and glycerol—while TAGs and CEs, the least polar species, are consumed (Supplementary Fig. S23). The polarity increase thus reflects ongoing catabolism. In contrast, oleate supplementation lowers polarity because excess FFAs are re-esterified into TAG and CE, as also indicated by increased LD volume^50^. Cancer-derived HeLa cells likely depend more strongly on lipid catabolism, explaining their consistently higher LD polarity. By contrast, differentiated 3T3-L1 adipocytes, specialized for lipid storage and enriched in TAG and CE, contained the largest LDs but displayed the lowest polarity, even under starvation (Fig. 3n, Supplementary Fig. S24). This polarity is even closely matching that of pure TAG and CE (Supplementary Fig. S8). This underscores that adipocytes prioritize stable energy storage, maintaining strict control over lipid mobilization in contrast to the flexible response of transformed cells. Together, these findings show that LD polarity heterogeneity reflects metabolic state and lipid flux. The rise in polarity with catabolism points to ongoing ester hydrolysis and FFA release, suggesting that LD polarity may serve as a dynamic readout of lipolysis and its coordination with other organelles.

### ExSPM unveils high-polarity LDs as a hallmark of lysosome-dependent acid lipolysis (lipophagy)

Lipolysis^51^, a primary pathway for LD breakdown, occurs via cytoplasmic neutral lipolysis or acid lipolysis, the latter also known as lipophagy^49, 51^. In lipophagy, autophagosomes engulf LDs and fuse with lysosomes, where acid lipase hydrolyzes TAG and CE to release FFAs. Lysosomes may also directly contact LDs to mediate lipid transfer and hydrolysis^52^. Using ExSPM with Nile Red, followed by fixation and LAMP1 immunostaining, we found that in normal COS-7 cells, high-polarity LDs strongly colocalized with lysosomes, whereas low-polarity LDs did not (red arrows in Supplementary Fig. S25). However, under OA treatment, high-polarity LDs were nearly absent, and low-polarity LDs were largely excluded from lysosomes. To confirm that high-polarity LDs were true LDs rather than Nile Red–labeled lysosomes, we performed colocalization with LS488—a validated LD-specific dye (Fig. 1)—and lysosomal markers (Fig. 4a). In normal cells, LS488-only LDs were predominantly low in polarity (cyan arrows in Fig. 4b and Supplementary Fig. S26), while LDs positive for both LS488 and lysosomes displayed higher polarity with reduced LS488 brightness (orange arrows). Lysosome-only LDs exhibited the highest polarity and lacked LS488 signal (red arrows). Quantitative analysis revealed a stepwise polarity increase after lysosomal fusion (Fig. 4c). These results may suggest a model in which neutral lipids undergo hydrolysis, leading to increased LD polarity and progressive loss of LS488 signal, consistent with earlier LS488–LiveDrop data (Fig. 1). Starvation further increased lysosome abundance and enhanced lipophagy, resulting in more lysosome-associated high-polarity LDs (Fig. 4d-4f and Supplementary Fig. S27), further reinforcing the model of polarity increase during lipophagy.

**Figure 4.**
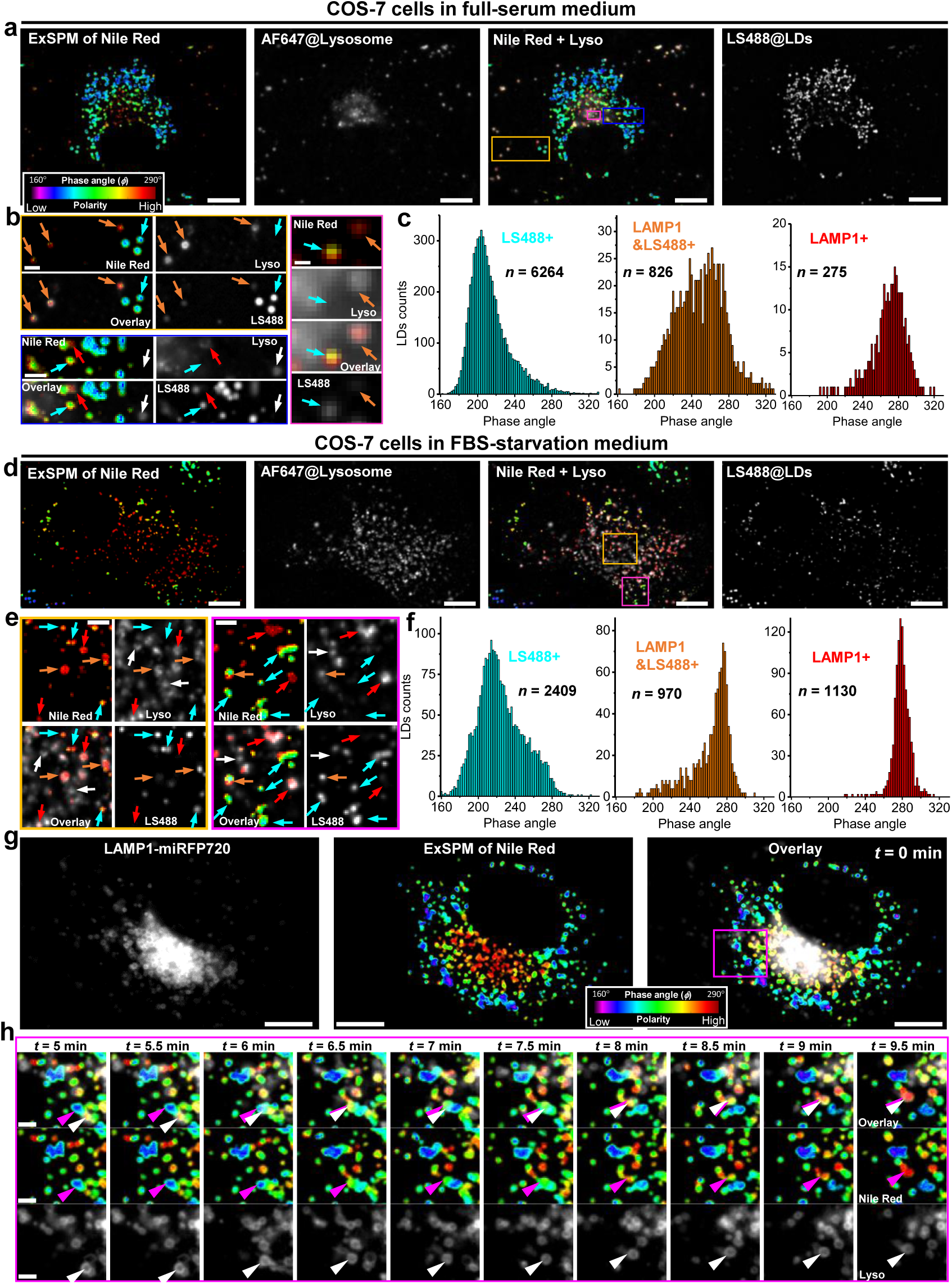
ExSPM unveils high-polarity LDs as a hallmark of lysosome-dependent acid lipolysis. **a.** Concurrent imaging of a COS-7 cell in full-serum medium, showing the ExSPM-derived LD polarity map (Nile Red staining), AF647-labeled lysosomes, and LipidSpot488 (LS488)-labeled LDs. **b.** Zoomed-in views of the boxed regions in a. Cyan arrows: LS488-only LDs; Orange arrows: LDs labeled by both LS488 and lysosomes; Red arrows: lysosome-only LDs; White arrows: free lysosomes not associated with LDs. **c.** Distribution of phase angles (polarity) for individual LDs across 30 cells from three independent experiments, categorized by different markers (LS488, lysosomes, or both). **d.** Concurrent imaging of a COS-7 cell cultured in FBS-starvation medium for 16 h, showing the ExSPM-derived LD polarity map (Nile Red staining), AF647-labeled lysosomes, and LipidSpot488 (LS488)-labeled LDs. **e.** Zoomed-in views of the boxed regions in d. Yellow-boxed region, rotated 90° clockwise for clarity. Cyan arrows: LS488-only LDs; Orange arrows: LDs labeled by both LS488 and lysosomes; Red arrows: lysosome-only LDs; White arrows: free lysosomes not associated with LDs. Both free lysosomes and lysosomes associated with LDs significantly increased compared to these in a and b. **f.** Distribution of phase angles (polarity) for individual LDs across 20 cells from three independent experiments, categorized by different markers (LS488, lysosomes, or both) as c. **g.** Live-cell concurrent imaging of a COS-7 cell cultured in FBS-starvation medium for 2 h, showing the ExSPM-derived LD polarity map (Nile Red staining) and miRFP720-labeled lysosomes at *t*=0 min. **h**. Time sequences for the magenta boxed region in g, where a low-polarity LD migrated toward and fused with a lysosome, rapidly shifting to high polarity before disappearing. Magenta arrows indicate LDs, and white arrows indicate lysosomes. Scale bars: 10 μm (a, d, g); 2 μm (b, e, h); 0.5 μm for magnified view (magenta boxed region in b).

To directly track polarity changes, we performed time-lapse live-cell imaging of ExSPM-Nile Red together with spectrally separated miRFP720-labeled lysosomes. In starved COS-7 cells, a low-polarity LD moved toward and fused with a lysosome. After fusion, a portion of neutral lipids entered the lysosome, where they rapidly shifted to high polarity before disappearing, providing direct evidence that polarity increases reflect lysosome-mediated hydrolysis (Fig. 4g-h; Supplementary Videos 7). Notably, the complete transition—from low polarity to high polarity and eventual signal loss— occurred within ∼5 min, underscoring the efficiency of lysosomal lipid breakdown in meeting acute energy demands. Interestingly, some high-polarity LDs did not colocalize with lysosomes, suggesting that alternative, lysosome-independent pathways may also contribute under nutrient stress.

### ExSPM unveils high-polarity LDs as a hallmark of neutral lipolysis through LD-mitochondria contacts

The discovery of adipose triglyceride lipase (ATGL) established cytoplasmic neutral lipolysis as a major pathway for triacylglycerol hydrolysis, producing diacylglycerols and FFAs^51^. This process depends on LD– mitochondria proximity, enabling direct FFA transfer for β-oxidation^49, 53, 54^. To examine whether such contacts influence LD polarity, we imaged Nile Red–labeled LDs alongside mitochondria and LS488 in the same cells (Fig. 5a-d). In normal conditions, most LDs without mitochondrial contact were low in polarity, whereas a subset of mitochondria-associated LDs displayed higher polarity (Supplementary Fig. S28). Under FBS starvation, this effect intensified, with high-polarity LDs frequently enriched at LD–mitochondria interfaces (orange and red arrows in Fig. 5e and Supplementary Fig. S29). In contrast, low-polarity LDs rarely engaged mitochondria under starvation (cyan arrows in Fig. 5e), and some high-polarity LDs appeared without mitochondrial contact (white arrows in Fig. 5f), consistent with lysosome-mediated lipophagy as an alternative pathway. Quantitative analysis confirmed a strong correlation between LD–mitochondria association and polarity increase (Fig. 5g).

**Figure 5.**
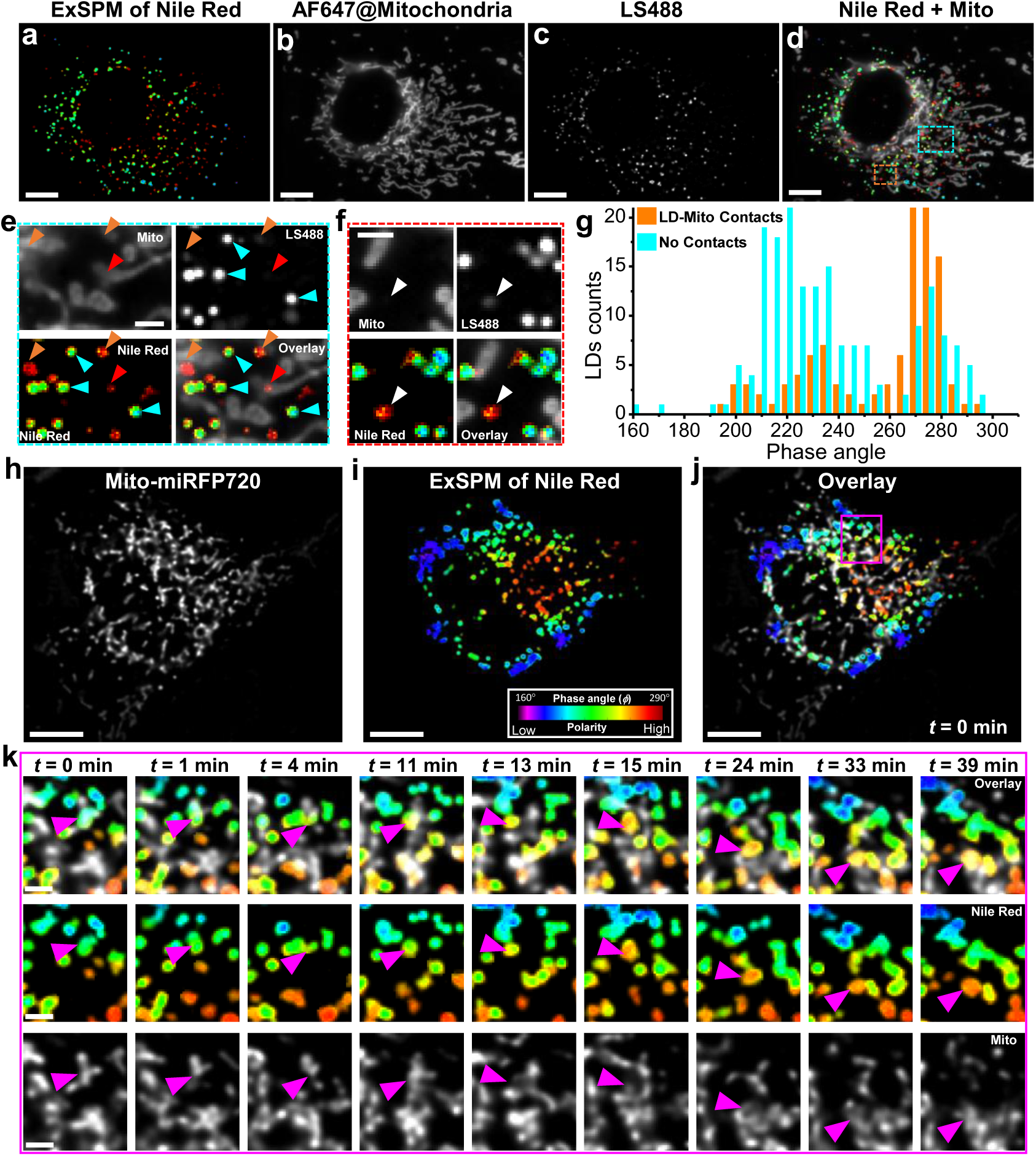
ExSPM unveils high-polarity LDs as a hallmark of neutral lipolysis through LD-mitochondria contacts. **a-d.** Concurrent imaging of a COS-7 cell in FBS-starvation medium, showing the ExSPM-derived LD polarity map (Nile Red staining) (a), AF647-labeled mitochondria (b), LipidSpot488 (LS488)-labeled LDs (c), and the overlay of polarity map and mitochondria (d). **e.** Zoomed-in views of the cyan boxed regions in d. Cyan arrowheads: LS488-only LDs without mitochondria contacts. Orange arrowheads: LDs labeled by LS488 in contact with mitochondria. Red arrowheads: LDs in contact with mitochondria but without LS488 labeling. **f.** Zoomed-in views of the orange boxed regions in d. White arrowheads: LS488-only LDs without mitochondria contacts but exhibiting high polarity. **g.** Distribution of phase angles (polarity) for individual LDs in the cell, categorized by the presence or absence of mitochondria contacts. **h-j.** Live-cell concurrent imaging of a COS-7 cell cultured in FBS-starvation medium for 2 h, showing miRFP720-labeled mitochondria (h), the ExSPM-derived LD polarity map (Nile Red staining) (i), and their overlay (j) at *t*=0 min. **k**. Time-lapse sequences of the magenta boxed region in (j), showing a low-polarity LD (magenta arrows) in stable contact with mitochondria that gradually shifted to higher polarity over ∼40 min. Scale bars: 10 μm (a-d, h-j); 2 μm (e, f, k).

To further resolve these dynamics, we performed time-lapse live-cell imaging of ExSPM-Nile Red and miRFP720-tagged mitochondria. In starved COS-7 cells, low-polarity LDs in stable contact with mitochondria gradually shifted to higher polarity over ∼40 min (magenta arrows in Fig. 5h–k, Supplementary Fig. S30 and Supplementary Videos 8–9). This was markedly slower than the ∼5 min polarity shifts observed during lysosomal fusion. Such gradual changes may reflect liquid-crystalline phase transitions within LDs^20^, driven by TAG consumption at LD–mitochondria interfaces during starvation. Supporting this, OA-treated cells—with predominantly low-polarity LDs—showed minimal LD–mitochondria association (Supplementary Fig. S31). Together, these findings identify neutral lipolysis at LD–mitochondria contacts as a key driver of polarity remodeling, operating in parallel with lysosome-mediated lipophagy. Both pathways appear to be co-regulated by metabolic factors such as mTOR, AMPK, and FOXO^51^, functioning as interconnected lipid catabolic processes that collectively sustain energy homeostasis.

### ExSPM quantifies lipolytic pathways and reveals LD catabolism is not governed by independent routes but by a tightly interconnected regulatory network

While both pathways contribute to lipid catabolism, it is not surprising that cells can dynamically regulate their usage for LD hydrolysis depending on metabolic conditions. To examine this regulation, we performed concurrent imaging of ExSPM-Nile Red, lysosomes, and mitochondria, defining high-polarity LDs (phase angle > 240°) as actively lipolytic (Fig. 6a–b). Lipolytic LDs fell into four categories: contacting lysosomes only, both lysosomes and mitochondria, mitochondria only, or neither. These broadly correspond to acid lipophagy (first two) and neutral lipolysis (last two).

**Figure 6.**
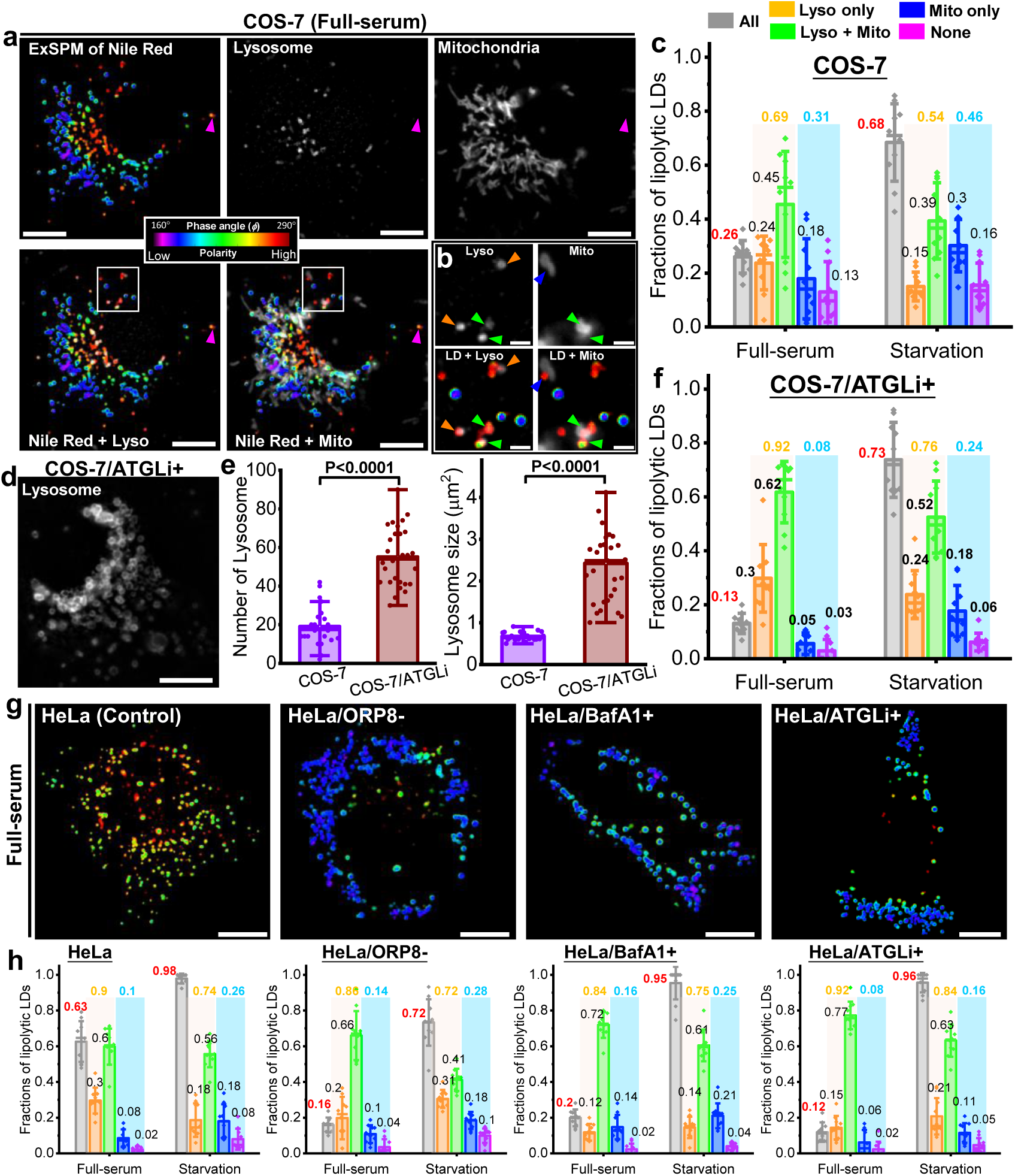
ExSPM quantifies lipolytic pathways and reveals a tightly interconnected regulatory network for LD catabolism. **a.** Concurrent imaging of a COS-7 cell in full-serum medium, showing the ExSPM-derived LD polarity map (Nile Red staining), AF647-labeled lysosomes, AF488-labeled mitochondria, and the overlay of polarity map and organelles. **b.** Zoomed-in views of the boxed region in a. Orange arrows: lysosome-only LDs; Green arrows: LDs contacting both mitochondria and lysosomes; Blue arrows: mitochondria-only LDs; Magenta arrows: free LDs not associated with either organelle. **c.** Statistical analysis of fractions of lipolytic LDs in COS-7 cells under nutrient-rich and deprivation conditions, each derived from analyses across 10 cells from three independent experiments. The gray bar represents the overall lipolysis ratio, referring to the proportion of lipolytic LDs relative to the total LDs in a single cell. The orange, green, blue, and magenta bars represent the proportions of lipolytic LDs contacting lysosomes only, both lysosomes and mitochondria, mitochondria only, or neither, respectively, within the total lipolytic LDs of a single cell. The first two categories are classified as acid lipophagy, while the latter two are classified as neutral lipolysis. **d.** A representative image of lysosomes (anti-LAMP1) in an Atglistatin-treated COS-7 cell (COS-7/ATGLi+), showing a marked increase in both the number and volume of lysosomes. **e.** Quantification of lysosome number and size in normal COS-7 and COS-7/ATGLi+ cells. Each dataset was obtained from at least 24 cells from three independent experiments. Two-tailed Student’s t-tests were used to determine *P* values. **f.** Statistical analysis of fractions of lipolytic LDs in ATGLi-treated COS-7 cells under nutrient-rich and deprivation conditions, each derived from analyses across 10 cells from three independent experiments. **g.** Representative ExSPM-derived LD polarity maps of a normal HeLa cell, ORP8-knockout HeLa cell (HeLa/ORP8−), bafilomycin A1-treated HeLa cell (HeLa/BafA1+), and Atglistatin-treated HeLa cell (HeLa/ATGLi+). **h.** Quantification of lipolytic LD fractions in HeLa, HeLa/ORP8−, HeLa/BafA1+, and HeLa/ATGLi+ cells under nutrient-rich and deprivation conditions, based on analyses of 10 cells from three independent experiments. In c, e, f and h, error bars represent mean ± s.d., and each point represents a single cell. Scale bars: 10 μm (a, d, g); 2 μm (b).

In COS-7 cells under nutrient-rich conditions, ∼26% of LDs were actively lipolytic, with lipophagy accounting for the majority (∼69%; Fig. 6c, left). Strikingly, most of these lipophagic LDs also maintained mitochondrial contact, suggesting that lysosome– mitochondria cooperation, rather than lysosome-only activity, plays the predominant role in basal lipid turnover. Neutral lipolysis contributed to ∼31% of breakdown events. Inhibition of ATGL with Atglistatin (ATGLi)^55^, the key cytosolic lipase, reduced neutral lipolysis to 8% and halved overall lipolysis to 13% (Fig. 6d–f, Supplementary Fig. S32), despite a pronounced expansion of lysosomes in both number and volume (Supplementary Fig. S7). This thus indicates that although lysosomal capacity increases, lipophagy cannot fully compensate for the loss of neutral lipolysis. Inversely, under acute nutrient deprivation, lipolysis increased substantially in both COS-7 and COS-7/ATGLi+ cells, with acid lipophagy and neutral lipolysis synergistically upregulated to meet energy demands (Fig. 6c and 6f, right). However, prolonged starvation shifted the balance toward neutral lipolysis, challenging the prevailing view that chronic stress universally favors autophagy. These results show that ExSPM can not only quantify the extent but also the pathway-specific kinetics of LD catabolism.

Compared with COS-7, HeLa cells showed far stronger lipid catabolism, with 63% of LDs actively lipolytic in full serum and up to 98% under starvation (Fig. 6g-h, left), with a stronger reliance on lipophagy (90% and 74%, respectively). To dissect pathway dependence, we perturbed lipophagy and neutral lipolysis using ORP8 knockout^56^, bafilomycin A1 (BafA1)^57^, or ATGLi (Fig. 6g-h, Supplementary Fig. S33). ORP8 loss disrupts selective lipophagy, where LDs are specifically targeted to autophagosomes, whereas BafA1 prevents lysosomal acidification, broadly impairing general autophagy. Both manipulations markedly reduced basal lipolysis to ∼20%, underscoring the central role of lysosomes. Intriguingly, neutral lipolysis did not compensate in either condition, implying that cytosolic lipase activity also depends on lysosomal integrity—possibly through regulatory mechanisms such as surface protein degradation or mTOR/AMPK signaling. Moreover, lysosome-only contacts declined, while LDs simultaneously contacting lysosomes and mitochondria increased, suggesting a compensatory mechanism in which mitochondria sustain lipid breakdown when canonical lipophagy is impaired. In contrast to COS-7 cells, ATGL inhibition only mildly reduced the relative contribution of neutral lipolysis in HeLa cells, consistent with its minor role in basal lipolysis. Strikingly, however, ATGLi drastically dropped overall lipolysis to 12%, suggesting that ATGL inhibition also impairs lipophagy.

As in COS-7, nutrient deprivation strongly stimulated lipolysis in HeLa cells, even when individual pathways were suppressed. ORP8 knockout had the most pronounced effect, limiting lipolysis to ∼72%, whereas BafA1 and ATGL inhibition still permitted ≥95% (Fig. 6h). This resilience highlights the redundant metabolic capability of cells to meet the acute energy demand. Under starvation, neutral lipolysis became more prominent again, likely because lysosomes were heavily engaged in other autophagic processes such as mitophagy and ribophagy, reducing their availability for lipophagy. In this context, cytosolic lipases—less constrained by lysosomal capacity—were therefore preferentially mobilized to maintain free fatty acid flux.

Together, these findings demonstrate the persistence of high lipolysis under acute nutrient deprivation despite pathway inhibition, reflecting compensatory redundancy that safeguards acute energy supply. Selective lipophagy, general autophagy, and neutral lipolysis thus operate as an integrated system, dynamically reallocating their contributions with nutrient status and lysosomal load, yet remaining coupled through a tightly interconnected regulatory network in which perturbation of one pathway reshapes the others.

## Discussion

ExSPM, integrating high-throughput excitation fluorescence spectral microscopy with phasor analysis, provides highly sensitive detection of subtle spectral shifts, enabling the visualization of intracellular heterogeneities at high spatial and temporal resolutions. By bypassing fitting-based spectral unmixing, spectral phasor analysis converts high-dimensional spectra into an intuitive two-dimensional representation, enabling direct visualization of small environmental differences without prior knowledge. Methodologically, it not only provides a highly sensitive tool but also redefines the capabilities of traditional fluorophores, potentially broadening the scope of biosensing for a wider range of cellular molecules and physicochemical parameters.

Biologically, ExSPM revealed unexpected richness in the polarity of LDs, overturning the traditional view of LDs as uniform inert depots. We observed marked heterogeneity in polarity across LDs within single cells, and even within individual LDs over time. High-polarity LDs consistently correlated with lipid catabolism, whereas low-polarity LDs were linked to lipid storage or buffering of excess lipids. These findings establish LD polarity as a functional marker of metabolic state rather than a passive physical property. Notably, polarity was not static: LDs undergoing interactions with lysosomes or mitochondria displayed progressive polarity increases, reflecting real-time hydrolysis of neutral lipids. This dynamic polarity shift at organelle contact sites provides direct evidence that acid lipophagy and neutral lipolysis act as localized, compartment-specific processes in lipid turnover. A central advance of this study lies in the dissection of pathway-specific contributions to LD catabolism. We found that, under basal conditions, lipophagy dominated LD turnover, while neutral lipolysis provided only minor contributions, particularly in cancer cells. However, under energy deprivation, neutral lipolysis was more strongly upregulated compared to lipophagy, likely due to lysosomal resource competition. This dynamic redistribution highlights the flexibility of lipid catabolic programs in adapting to cellular energy status.

Importantly, genetic and pharmacological perturbations revealed that inhibition of one pathway not only suppressed its direct contribution but also diminished the activity of others. For example, ORP8 knockout or lysosomal inhibition blunted both lipophagy and neutral lipolysis, while ATGL inhibition reduced overall catabolism beyond its expected contribution. These results suggest that the selective lipophagy, general autophagy/lipophagy, and cytosolic neutral lipolysis are tightly coupled through shared signaling hubs and interconnected regulatory network, rather than operating in a simple compensatory, “either–or” fashion. Such coupling may be mediated by global metabolic regulators such as AMPK and mTOR, or by cross-organelle communication involving lipid transporters, perilipins, or other cofactors. Thus, LD catabolism emerges as a systems-level property coordinated by an interdependent metabolic network.

The biological implications of this network architecture are profound. In HeLa cells, for instance, lipid catabolism was elevated compared with non-cancerous cells, with lipophagy playing a particularly prominent role under nutrient-replete conditions. Yet, even when major pathways were suppressed, high levels of lipolysis persisted during severe energy deprivation, reflecting compensatory redundancy that ensures continuous energy supply. Such flexibility may confer a selective advantage to cancer cells by enabling them to meet the fluctuating demands of growth and survival. More broadly, these findings imply that lipid catabolism is safeguarded by overlapping mechanisms, in which failure of one pathway reshapes rather than abolishes the network’s activity.

Future investigations into the molecular mechanisms governing LD polarity, as well as the multidirectional interplay between lipolytic pathways—such as interactions with lipases^58^, lipid transporters^59^, and organelles^2, 3^—could further refine our understanding of lipid catabolism. More broadly, viewing cellular lipid metabolism as an integrated network rather than isolated pathways should be key to uncovering the fundamental principles of lipid regulation. For example, because ORP8 mediates both selective LD engulfment by autophagosomes^56^ and LD biogenesis at ER–mitochondria contact sites^60^, its role in coordinating distinct lipolytic pathways warrants deeper exploration. Additionally, integrating ExSPM’s temporal resolution and AI-driven spectral analysis may enable automated, high-throughput metabolic monitoring^61^.

Together, our findings establish ExSPM as both a methodological advance and a conceptual framework for lipid metabolism, providing a powerful platform to dissect intracellular heterogeneity, metabolic regulation, and organelle communication with broad implications for energy homeostasis, cancer metabolism, and lipid-associated diseases.

## Acknowledgments

This work was supported by the National Natural Science Foundation of China (62205048, 62475032).

## Author Contributions

K.C. conceived the research. J.L. designed and conducted the experiments. All authors contributed to experimental designs, data analysis, and paper writing.

